# Spatial confinement modulates macrophage response in microporous annealed particle (MAP) scaffolds

**DOI:** 10.1101/2022.08.16.504156

**Authors:** Yining Liu, Alejandra Suarez-Arnedo, Lindsay Riley, Tasman Miley, Jingyi Xia, Tatiana Segura

## Abstract

Macrophages are essential in the initiation, maintenance, and transition of inflammatory processes like foreign body response and wound healing. Mounting evidence suggests that physical factors also modulate macrophage activation *in vitro* and *in vivo*. 2D *in vitro* systems have demonstrated that constraining macrophages to small areas or channels modulates their phenotypes and changes their responses to known inflammatory agents such as lipopolysaccharide. However, how dimensionality and pore size affect macrophage phenotype is less explored. In this work, we studied the change in M1/M2 polarization when macrophages were confined in microporous annealed particle scaffolds (MAP), which are granular hydrogels generated from annealed spherical microgels. We engineered three types of MAP gels comprising 40, 70, and 130 µm diameter particle sizes, respectively. Particles sizes were selected using outputs from software LOVAMAP that analyzes the characteristics of 3-D pores in MAP gels. Since the size of building block particle correlates with pore size inside the final scaffolds, our three scaffold types allowed us to study how the degree of spatial confinement modulated the behavior of embedded macrophages. Spatially confining macrophages in scaffolds with pore size on the scale of cells led to a reduced level of the inflammatory response, which was correlated with a change in cell morphology and motility.

## Introduction

Macrophages are central to many injuries and diseases^1^. During a typical episode of inflammation, macrophages are among the first responders to arrive and polarize into a variety of activation states to perform specific functions. These states can be simplified as a spectrum from pro-inflammatory (M1) to pro-reparative (M2) phenotypes^2, 3^. Generally, M1 phenotypes are associated with the initiation and maintenance of inflammation while M2 phenotypes are closely tied to the resolution of inflammation and the changeover to the regeneration phase^4^. Besides the intrinsic differentiation pathways governing this timely transition in phenotypes, macrophages are also adaptive to microenvironmental cues from neighboring cells and the extracellular matrix in which they reside^5^. Biochemical factors secreted by other cells, like IFN-γ or IL-4, can direct macrophages into pro-inflammatory or pro-reparative phenotypes^6^. The molecular mechanisms behind these common soluble factors and their effects on macrophages have been extensively studied. However, the mechanisms by which physical signals regulate macrophage activation are less explored^7–9^. In the field of biomaterials, researchers have tested the influences of a wide range of material properties on macrophage modulation in pursuit of better biocompatibility. For example, surface modification by increasing hydrophilicity reduces macrophage attachment, while decorating surface with cell-binding ligands biases macrophage polarization^10–13^. Understanding the specific mechanotransduction mechanisms that govern phenotypic macrophage changes will guide future biomaterial design and achieve a far-reaching physiological significance.

Spatial confinement is a well-known parameter for regulating macrophage response in the context of tissues or material scaffolds. Topographical designs that force macrophages into an elongated cell shape have been shown to promote a pro-regenerative M2 phenotype^14^. By using micropatterned surfaces, microporous substrates, and cell crowding to induce spatial confinement, investigators were able to prevent mouse bone marrow-derived macrophages or RAW264.7 cells from spreading, thereby suppressing late lipopolysaccharide (LPS)-related transcriptional programs and cytokine expression^15^. Actin polymerization was limited in macrophages within a confined space, which reduced nuclear translocation of the actin-dependent transcription co-factor, myocardin-related transcription factor-A^15^. This factor regulates the LPS-stimulated inflammatory response, and its reduction in macrophages results in a lower phagocytic potential and less pro-inflammatory cytokine secretion^15^. Macrophages undergoing spatial constraints also experience chromatin compaction and epigenetic alterations (e.g., reduced histone deacetylase 3 level and enhanced H3K36-dimethylation)^16^.These findings revealed some key pathways that guide macrophage response to spatial confinement. In another study, a precisely-defined fiber poly(ε-caprolactone) scaffold with 40 μm wide box-shaped pores was shown to facilitate cell elongation in primary human macrophages when compared to larger pores, driving an M2-like polarization with increased M2 gene expression (CD206, CD163, IL-10) and reduced M1 activity (inflammatory cytokine expression and phagocytic activity)^17^. Together, these studies support the notion that the modulation of macrophage cell shape by spatial confinement impacts macrophage response^18^.

In the past decade, granular materials have emerged as an option for well-defined *in vitro* systems with plug-and-play components and tissue-mimicking 3D environments^19–22^. When co-culturing hMSCs with M1 macrophages under pro-inflammatory conditions (i.e., conditioned medium from M1 macrophages) in packed granular microgels^23^, IL-10 was identified as a key modulating factor. Immobilization of IL-10 onto microgels was developed as a method for controlling macrophage polarization in 3D culture. Another study showed that microgel size in granular gels modulated murine macrophages cell line polarization^24^. Specifically, large pore size promoted a more M2 phenotype, and small pore size induced a more M1 phenotype, both *in vitro* and *in vivo*^24^. However, these results are not in line with previous findings observed in primary murine macrophages that spatially confinement macrophages reduced their pro-inflammatory response. In this study, we used microporous annealed particle (MAP) scaffolds, a granular gel formed by interlinking packed microgels as our 3D culture system, and we explored the response of bone marrow-derived macrophages (BMDM) within MAP scaffolds comprising 40 μm, 70 μm, and 130 μm diameter microgels. We find that macrophage activation levels correlated with changes in morphology, cell motility, and nucleus shape, which were regulated by the pore sizes of MAP scaffolds.

## Results

### LOVAMAP guided the design of MAP scaffolds with controllable 3D-pore size

To create MAP scaffolds with the desired degree of spatial confinement for our target cell type (BMDM), we used a computational approach to guide our design. We started by simulating an array of monodisperse MAP scaffolds with varying microgel size (from 40 µm to 200 µm) using SideFX Houdini software, then extracted 3D-pore data using LOVAMAP^25^, an in-house analytical software for granular materials (Figure 1a,b). 3D-pore volume was used as an approximation for the degree of spatial confinement since pore volume directly impacted encapsulated cell volume inside the scaffold. The prior 2D and 2.5D studies of BMDM macrophage confinement reported that macrophage confinement occurred in a space less than 4.2 pL in volume^14, 15^. Therefore, we chose scaffolds made with 40 µm diameter particles as the most confined condition since their median 3D-pore volume (3.2 pL) was below 4.2 pL, smaller than untreated BMDM (17 pL), and similar to reported *in vivo* alveolar (AV) murine macrophage volume^26, 27^ (Figure 1c). MAP scaffolds comprising 130 µm diameter particles were selected as an unconfined condition because they had a medium pore volume (96.2 pL) almost double the size of LPS-activated BMDM (51 pL) (Figure 1c). These 130 µm particles were also the largest size that could fit through the 29 and 31 Gauge syringe needles for *in vivo* scaffold injections without deformation^28^. In addition, we opted for a less confined group using MAP scaffolds with 70 µm diameter particles, which had a median pore volume (15.8 pL) that matched untreated BMDM (Figure 1c).

**Figure 1.**
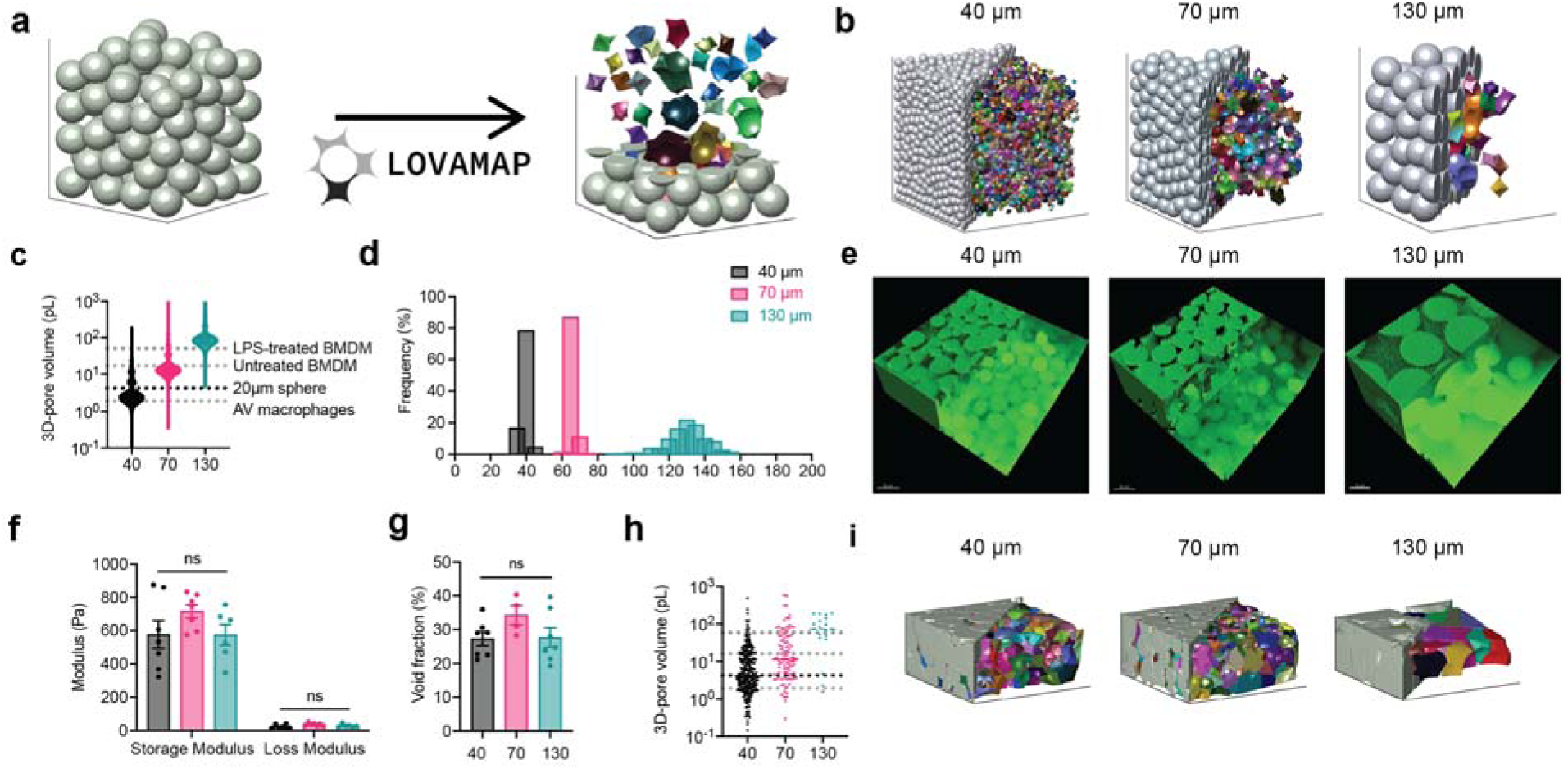
LOVAMAP software guided the design of MAP GELS with controllable 3D-pore size. a, LOVAMAP processes the images of MAP scaffolds and extracts information regarding 3D-pores inside the scaffolds. Microgels are grey and 3D-pores are colored, shown separated from one another to highlight variation in form. **b,** LOVAMAP outputs of three simulated MAP GELS comprising 40 µm, 70 µm and 130 µm diameter spherical microgels. **c,** 3D-pore volumes for 40 µm, 70 µm and 130 µm diameter simulated MAP scaffolds (n = 10 scaffolds per type) compared to the size of previously reported alveolar (AV) macrophages, untreated BMDM, and LPS-treated BMDM**. d,** Microgels were monodisperse in size (40 ± 2.9 μm, 70 ± 3.3 μm, and 130.8 ± 9.2 μm). **e,** Fluorescent images showing 40 µm, 70 µm and 130 µm diameter MAP scaffolds (scale bar 50 µm). **f,** Three types of MAP scaffolds had comparable storage and loss modulus. **g,** Void fraction in MAP scaffolds. Void fraction = (Total volume - Microgel volume) / Total volume. **h,** 3D-pore volumes for 40 µm, 70 µm and 130 µm diameter of lab-derived MAP scaffolds **i,** LOVAMAP outputs of lab-derived 40 µm, 70 µm and 130 µm MAP scaffolds. Microgels are grey, and 3D-pores are colored. Statistical analysis: one-way ANOVA with Tukey’s multiple comparisons test made between 40 µm, 70 µm and 130 µm MAP scaffolds groups only when there was significant difference among means. * p<0.05, ** p<0.01, *** p<0.001, **** p<0.0001. Error bars, mean ± s.e.m. n = 6 samples per scaffold type.

Using microfluidic devices, we generated microgels with an average diameter of 40 µm, 70 µm, and 130 µm as previously discussed (Figure 1d)^29^. These microgels were made of identical chemical components and possessed similar mechanical properties. When annealed together to form MAP scaffolds, they produced scaffolds with similar void fractions (25-35%) but distinctly different internal landscapes^30^ (Figure 1e-g). Again, using LOVAMAP, we analyzed microscopic fluorescent images of lab-derived MAP scaffolds to estimate the actual 3D-pore parameters, which revealed similar values to simulated data (Figure 1h and i). The analysis of 3D-pore size distribution of the lab-derived MAP scaffolds is limited by confocal image acquisition and light scatter. Further, image boundaries result in clipped particles and pores that extend beyond the region of interest, which results in underestimation of pore size. Nevertheless, the median 3D-pore sizes were identified as 4.2, 11.5, and 71.8 pL for 40, 70, and 130 µm, respectively, which matched simulated data and showed that 3D-pore size in MAP scaffolds can be controlled by changing the microgel size.

### Macrophage morphology in MAP scaffolds correlated with particle size, not activation condition

When cultured on 2D surfaces, BMDMs elongate upon IL-4 (M2) activation, and exhibit pancake shape with large cell areas after LPS/INF-g (M1) activation^14, 15^ (Supplementary Figure 1a). Less is known regarding macrophage morphology change upon activation in 3D culture. In this study, we observed that upon M1(LPS/IFNγ) or M2(IL-4) activation, embedded macrophages in MAP scaffolds had similar cell morphology with unstimulated macrophages (Figure 2a). No statistical difference was observed between activation and no activation groups in terms of surface area, volume, ellipticity (oblate), ellipticity (prolate), and sphericity (Supplementary Figure 1b). However, a divergent size-dependent morphological change was observed when we combined the data per MAP scaffold type and studied the effect of MAP scaffold particle size on BMDM morphology (Figure 2b-g). Macrophages cultured in 130 µm MAP scaffold had larger cell surface area and volume within the less confined 3D-pore (Figure 2b, c). The surface area-to-volume ratio (SA/V), a measurement of the ability to exchange materials with the external environment^31^, remained the same across three size groups (Figure 2d).

**Figure 2.**
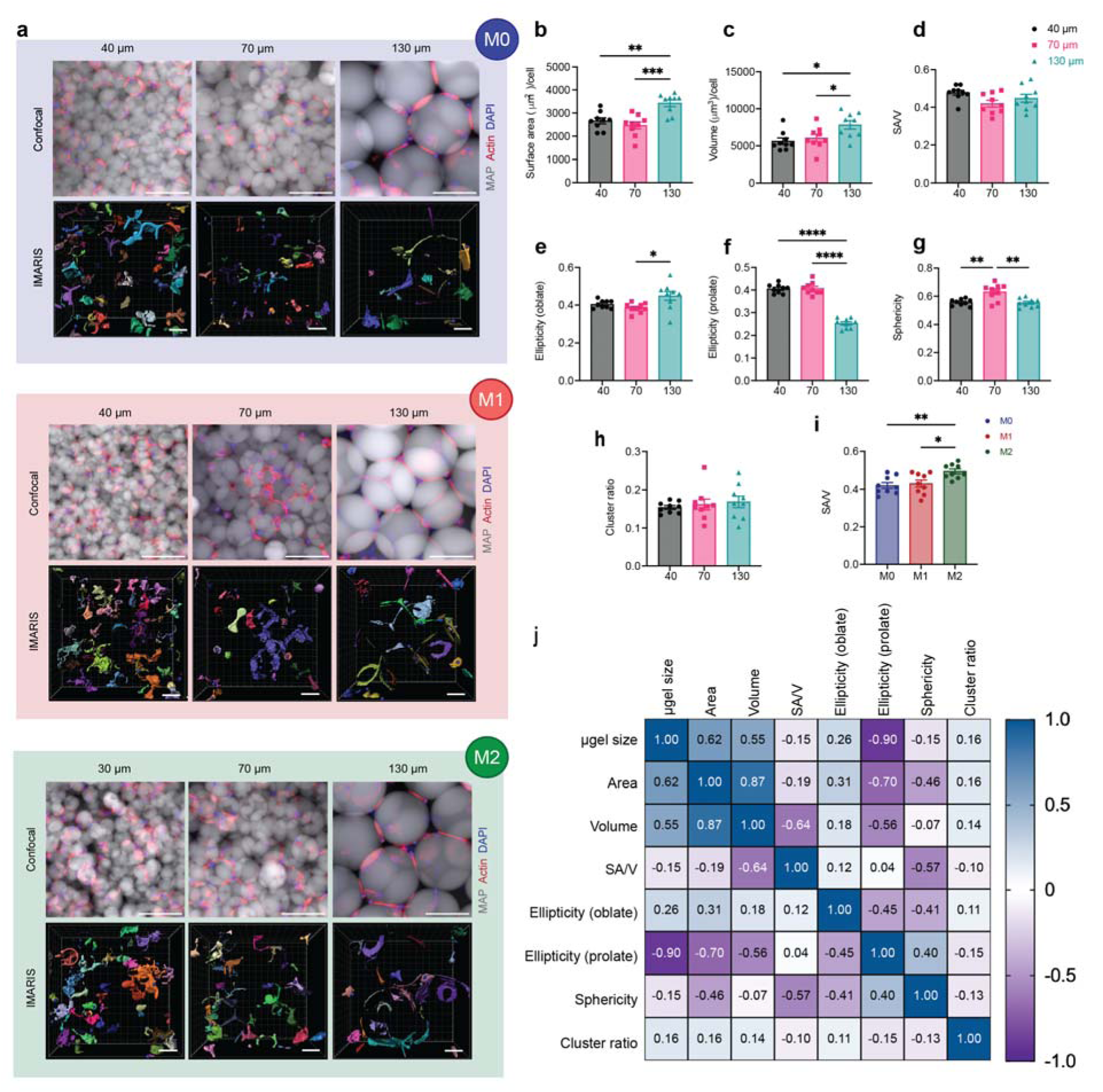
Macrophages showed divergent morphologies upon spatial confinement in MAP GELS. **a**, Fluorescent images showing BMDM encapsulated in MAP scaffolds with M0, M1 (LPS/IFNγ), and M2 (IL-4) activation (top row: microgels in grey, actin filaments in red, nuclei in blue, scale bar 100 μm) and the IMARIS software renderings of the single cells and cell clusters in MAP scaffolds (bottom row: colored units, scale bar 50 μm). **b-h**, Cell morphological parameters of BMDM encapsulated in different MAP scaffolds, combining three activation types: b, sum of total surface area normalized by cell number; c, sum of total volume normalized by cell number; d, surface area-to-volume ratio; e, ellipticity (oblate); f, ellipticity (prolate); g, sphericity; h, cluster ratio. **i,** Surface area-to-volume ratio of cells cultured in MAP scaffolds after M0, M1, and M2 activation. **j**, Correlation matrix of cell morphological parameters (blue represents positive correlation and purple represents negative correlation). Statistical analysis: one-way ANOVA with Tukey’s multiple comparisons test made between 40 µm, 70 µm and 130 µm MAP scaffolds groups only when there was significant difference among means. * p<0.05, ** p<0.01, *** p<0.001, **** p<0.0001. Error bars, mean ± s.e.m., n = 9 per group.

The ellipticity value characterizes the elongation of the cell/cluster by fitting an ellipsoid inside and calculating the degree of deviation from circularity. The ellipticity prolate and oblate represents the directionality of the longer axis of the ellipsoid, with prolate ellipsoid lengthening vertically and oblate ellipsoid flattening horizontally (Supplementary figure 1b). A significantly lower ellipticity (prolate) and a higher ellipticity (oblate) was observed in 130 µm MAP scaffolds (Figure 2f,e). This observation suggested that BMDM took a more elongated cell shape in the vertical direction, possibly due to nutrient gradients, 3D-pore topography, and gravity. The sphericity of cells in 40 µm MAP scaffolds and 130 µm MAP scaffolds was drastically lower than in 70 µm MAP scaffolds (Figure 2g), again pointing to a less spherical, more rough-edged morphology. Further BMDMs formed cell clusters to the same degree in all 3D-pores sizes, indicating that cell movement was not obstructed (Figure 2h).

We next aggregated the data per cytokine activation type, rather than scaffold size. We found that the only variable that presented statistical difference upon macrophage activation was SA/V, showing higher values upon M2 activation (Figure 2i). In the context of macrophages, higher SA/V had been associated with M1 activation since an increased surface area per cellular volume allows the cell to be more efficient in pathogen detection and phagocytosis^32, 33^. However, the tendency of a lower volume per cell upon M2 activation (Supplementary Figure 1c) led to a significant change in SA/V even though the surface area per cell remained constant upon both M1/M2 activation in contrast with unstimulated macrophages. Thus, this data again showed that the biggest factor affecting macrophage morphology in MAP scaffolds is microgel size.

Last, we used a Pearson’s correlation matrix to further tear out the interactions between all variables we collected and analyzed (Figure 2j). Both surface area and volume positively correlated with microgel size, while ellipticity prolate had a negative correlation with microgel size. Sphericity also negatively correlated with SA/V, as reported previously^34^. Macrophages cultured in 130 µm MAP scaffolds with less spatial confinement were able to elongate and spread out, while macrophages cultured in 70 µm MAP scaffolds had higher cell sphericity and reduced SA/V ratio.

Collectively, the evidence showed that BMDMs changed their morphology as a function of the microgel size, which directly tied to the level of spatial confinement BMDM perceived within the scaffold.

### Macrophages showed divergent nuclear shape upon spatial confinement in MAP scaffolds

We looked at nuclear shape of encapsulated cells since it was connected to cell phenotype and activation state^14^. Previous studies have also demonstrated that changes in cell shape led to nuclear morphology changes and thereby causing the reorganization of nuclear components^14^. Without surprise, size-dependent change in cell morphology was mirrored in the change of nucleus shape (Figure 3 a, b). BMDMs in 130 µm MAP scaffolds had a significantly larger nuclei area, and both 40 µm and 130 µm MAP scaffolds elicited a less circular nuclei shape with a higher aspect ratio (Figure 3 c, d). The elevated cell elongation in 40 µm and 130 µm MAP scaffolds was associated with a more elongated nuclei shape, which was previously linked to a higher degree of chromatin condensation^35^ and histone acetylation^36^. Correlation matrix between cellular morphological parameters and nuclei shape revealed that there was a high correlation between cell size and nuclei size, meaning that bigger nuclei were observed in bigger cells. Analogous parameters of the nuclei, such as circularity, roundness, and solidity, had a highly positive correlation with the sphericity of the cells (Figure 3e and supplementary figure 2), which further demonstrated the positive interaction between cell morphology and nuclear shape.

**Figure 3.**
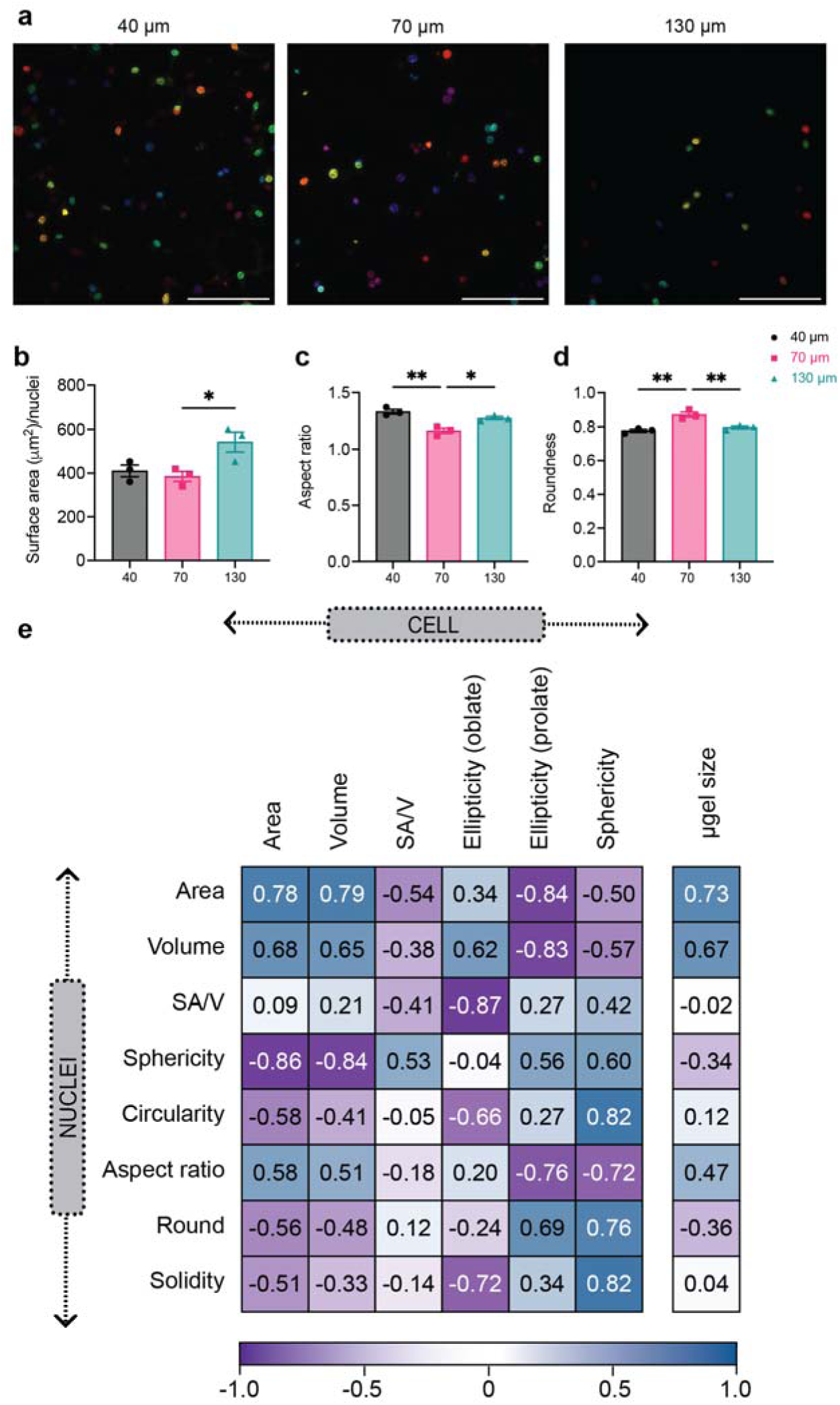
Macrophages showed divergent nuclear shape upon spatial confinement in MAP scaffolds. **a,** Mean intensity projection images of cell nuclei from confocal images of 3D MAP scaffolds scaffolds (scale bar 100 μm). **b-d**, morphology analysis of nuclei shape for unstimulated BMDM in MAP scaffolds, showing surface nuclei area, aspect ratio, and roundness. **e**, Correlation matrix of selected cellular morphology and nuclear shape characteristics for unstimulated BMDM in MAP scaffolds. Statistical analysis: one-way ANOVA with Tukey’s multiple comparisons test made between 40 µm, 70 µm, and 130 µm MAP scaffolds groups only when there was significant difference among means. * p<0.05, ** p<0.01, *** p<0.001, **** p<0.0001. Error bars, mean ± s.e.m., n = 3 from three independent experiments per scaffold type.

### Macrophage antigen presentation markers differed for LPS/IFNγ or IL-4 stimulation

We first wanted to confirm that macrophages cultured in 2D showed the expected differences when activated with M1(LPS/IFNγ) or M2(IL-4) using ELISA quantification of inflammatory cytokine secretion and flow cytometry analysis of inflammatory marker expression (Supplementary Figure 3a, b). The inflammatory phenotype was studied using well documented IL-6 and TNF cytokine secretion and iNOS and CD86 marker expression, and the regenerative phenotype was studied using Arg1 and CD206 maker expression. In addition, we added two markers of interest, MHCII and CD11c(antigen presenting markers), as macrophages are also recognized for their antigen presenting properties. As expected, M1(LPS/IFNγ) stimulation elevated the secretion of IL-6 and TNF and the expression of pro-inflammatory markers inducible nitric oxide synthase (iNOS) and CD86. We also found that the antigen-presenting marker MHCII was upregulated upon M1(LPS/IFNγ) stimulation compared to the unstimulated control and macrophages stimulated with M2(IL-4),indicating that M1 stimulation increased macrophage antigen presentation ability (Supplementary Figure 3c). On the other hand, stimulation of macrophages with M2(IL-4) did not induce the secretion of IL-6 and TNF but upregulated pro-regenerative markers CD206 and Arginase 1 (Arg1). Similar to M1(LPS/IFNγ) stimulation, we found that stimulation with M2(IL-4) increased macrophage antigen presenting ability, with an upregulation of CD11c (Supplementary Figure 3d).

### LPS/IFNγ-stimulated macrophages in MAP scaffolds reduced the inflammatory response

It’s known in the literature that forcing macrophages into elongated cell shape with topography design promoted a pro-regenerative M2 phenotype^14^, while spatially confining macrophages and thereby restricting cell spreading reduced their inflammatory response to LPS^15^. Thus, we next set out to study how spatially confined macrophages in MAP scaffolds respond to cytokine activation. Activating macrophages in MAP scaffolds with an M2(IL-4) stimulus or with no stimulus did not lead to a significant bias towards either pro-regenerative or pro-inflammatory markers (Supplementary figure 4). On the other hand, M1(LPS/IFNγ) activation led to a reduced macrophages inflammatory response, with a significant decrease in both pro-inflammatory marker expression (iNOS and CD86) as well as IL-6 and TNF cytokine expression in all MAP scaffolds groups (Figure 4a). These results indicated that confining BMDM in MAP scaffolds assembled with 40-130 µm microgels changed their inflammatory phenotypes. Specifically, iNOS and CD206 expression was significantly higher in 70 µm MAP scaffolds compared to 40 µm MAP scaffolds and 130 µm MAP scaffolds, pointing to a differential regulatory mechanism other than spatial confinement (Figure 4a, b). Remarkably, inflammatory cytokines TNF and IL-6 showed a 3D-pore size-regulated response (Figure 4a), with 40 µm MAP scaffolds drastically decreasing TNF and IL-6 secretion compared to 70 µm and 130 µm (Figure 4a). This observation aligned with previously reported results that spatial confinement reduced TNF and IL-6 production in M1(LPS) activated macrophages^15^.

**Figure 4.**
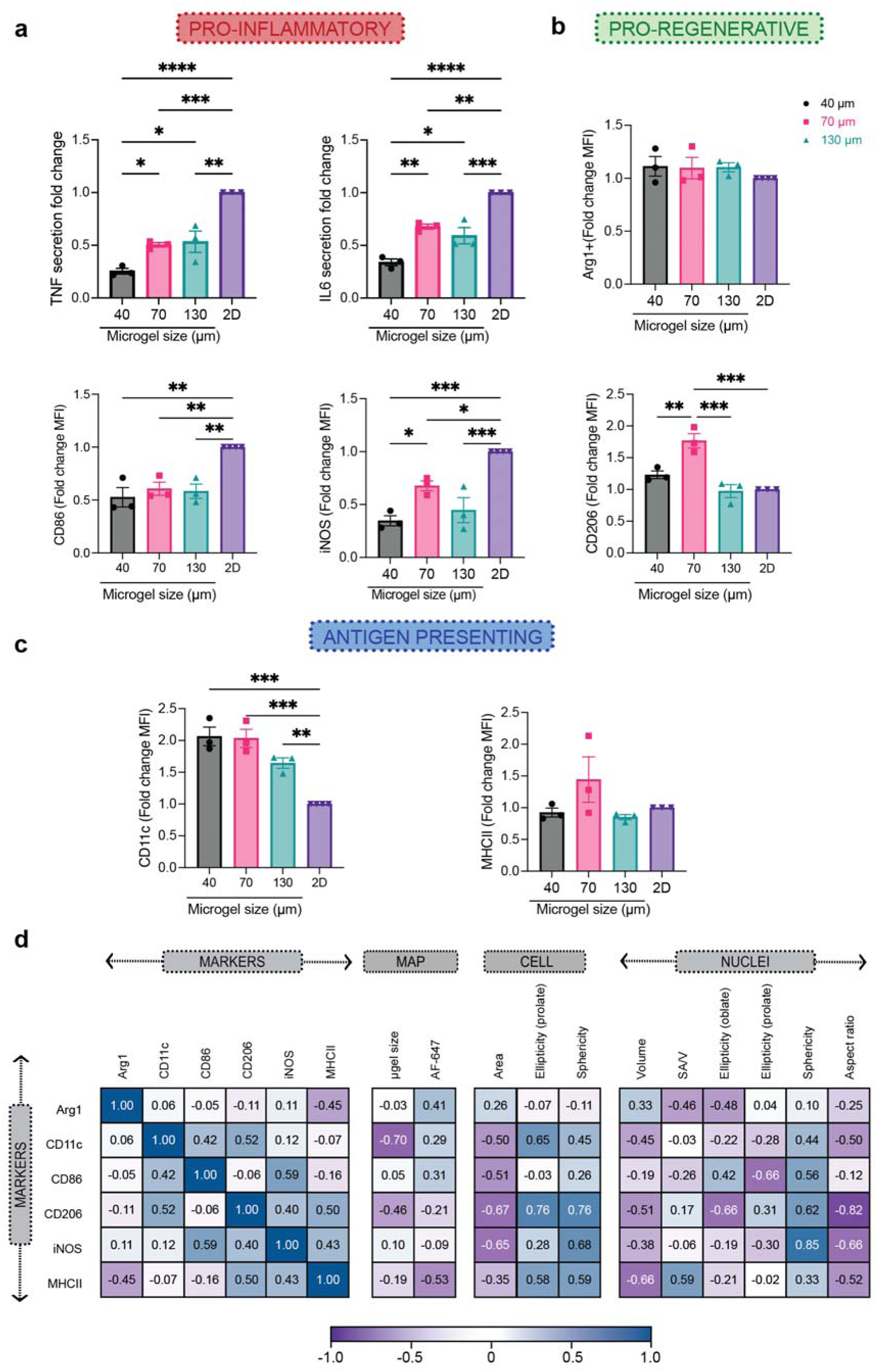
Macrophage phenotypes were governed by spatial confinement in MAP scaffolds upon M1(LPS/IFNγ) activation. **a,** Fold-change of inflammatory cytokine TNF and IL-6 secretion, and expression levels of pro-inflammatory markers iNOS and CD86, compared to the corresponding 2D culture with the same M1(LPS/IFNγ) activation. **b,** Expression levels of pro-regenerative markers CD206 and Arg1, compared to the corresponding 2D culture with the same M1(LPS/IFNγ) activation. **c,** Expression levels of antigen-presenting markers MHCII and CD11c compared to the corresponding 2D culture with the same M1(LPS/IFNγ) activation. **e**, Correlation matrix between phenotypical markers and selected morphological characteristics of BMDM encapsulated in MAP scaffolds upon M1(LPS/IFNγ) activation. Statistical analysis: one-way ANOVA with Tukey’s multiple comparisons test made between 40 µm, 70 µm and 130 µm MAP scaffolds groups only when there was significant difference among means. * p<0.05, ** p<0.01, *** p<0.001, **** p<0.0001. Error bars, mean ± s.e.m., n = 3 from three independent experiments per scaffold type.

### Antigen presentation for macrophages cultured in MAP scaffolds changed compared to 2D

In 2D culture experiment, MHCII was upregulated in macrophages with M1(LPS/IFNγ) stimulation. However, there was no difference in MHCII expression between BMDM cultured in the different MAP scaffolds groups, indicating that the 3D-pore size tested did not influence MHCII expression (Figure 4c). Though CD11c was upregulated for the IL-4 treatment group in 2D, we saw that its expression was higher for all BMDM cultured MAP scaffolds with M1(LPS/IFNγ) stimulation, and that the expression was higher than for MAP scaffolds treatment with IL-4. These results showed that the expression of CD11c in macrophages after cytokine stimulation dramatically changed for cells cultured in MAP scaffolds.

### Correlations between cellular morphology and cell phenotype were observed in MAP

Similar to the analysis we did to correlate cellular morphology and microgel size, we used a Pearson’s correlation matrix to uncover correlations between cellular morphology, MAP scaffold size and internalization, and macrophage phenotype with M1(LPS/IFNγ) stimulation (Figure 4d). Only CD11c showed a high negative correlation with the size of the microgels. The uptake of MAP (labeled with AF647) by infiltrated cells were observed^37^, but there is no high correlation between phenotypical markers and AF647 level. Interestingly, both the pro-inflammatory marker (iNOS) as well pro-regenerative marker (CD206) had a high positive correlation with the sphericity of cells/nucleus and a negative correlation with the area of a cell. This meant that smaller cells with a more spherical shape were associated with a higher expression of iNOS and CD206. The correlation between morphological characteristics and iNOS expression has not been reported previously. However, for CD206 the results obtained are contrary to what was observed in alveolar macrophages, where large cells express high MFI for CD206^38^. Similarly, CD86 showed a positive correlation with nuclear sphericity. Both pro-inflammatory cytokines are associated with nuclei circularity (Supplementary Figure 5). The clear associations between cellular morphology and cell phenotype aligned with the profiles in M1/M2 activation shown by previous publication, where macrophages with more circular shape presented a higher expression of M2-associated markers (CD206 and Arg1+)^24^. This observation lead us to suggest that differences in the association of microgel size with macrophages phenotype between this study and Lowen et al. work may be due to difference in packing on the microgels^39^ and cell size. Taken together, our results show that that the spatial confinement of MAP scaffolds modulates macrophages phenotype with M1(LPS/IFNγ) activation.

### Balance of M1/M2 phenotype upon M1 activation in MAP scaffolds

In traditional flow cytometry characterization of macrophage phenotypes, we used in this study, the macrophage population was first gated as CD11b+F4/80+ live cells, then further characterized by their expression levels of each functional marker. Median Fluorescent Intensity (MFI) was output to compare the expression of markers across samples. The basal levels of Arg1, CD206 and CD11c expression in BMDM were due to the differentiation method we used, which is known to induce a less inflammatory, pro-regenerative phenotype^40^ (Figure 5a). 2D cultured macrophage was used in each treatment condition as a reference to identify the main cell types associated with the treatments. Compared with the 2D group, macrophage cultured in MAP scaffolds M1(LPS/IFNγ) activation showed a reduction of double positive pro-inflammatory markers (iNOS+CD86+), mainly in 40 µm and 130 µm (Figure 5b,c) and a reduction of M1/M2 phenotype (CD206+iNOS+) in 40 µm MAP scaffolds (Figure 5d,e). These results were aligned with the increase in M1/M2 phenotype (Arg1+iNOS+) and double positive pro-regenerative markers (iNOS+CD86+) in 40 µm MAP scaffolds (Supplementary Figure 6a). The correlation matrix demonstrated a highly positive interaction between M1/M2 phenotype (CD206+iNOS+) and double positive pro-inflammatory markers (iNOS+CD86+). This association suggested that there was a macrophage subpopulation with high expression iNOS presenting both expressions of CD206 and CD86 (Figure 5f). There was a negative correlation between CD206+CD11c+ and CD206+CD86+ co-expression (Supplementary figure 6 and Figure 5f), showing that MAP scaffolds modulated macrophage response to M1 stimulus towards a more pro-regenerative, antigen-presenting phenotype. Similarly, high co-expresion CD206+MHCII+ showed a negative correlation with microgel size, and it was highly associated with the expression of CD11c+ and the ellipticity (prolate) of the cell (Figure 5f and Supplementary figure 6a). Altogether, macrophage phenotype landscapes in MAP scaffolds were shaped by 3D-pore size, and MAP scaffolds group showed distinctively lower responses towards M1 activation with a more pro-regenerative, antigen-presenting phenotype.

**Figure 5.**
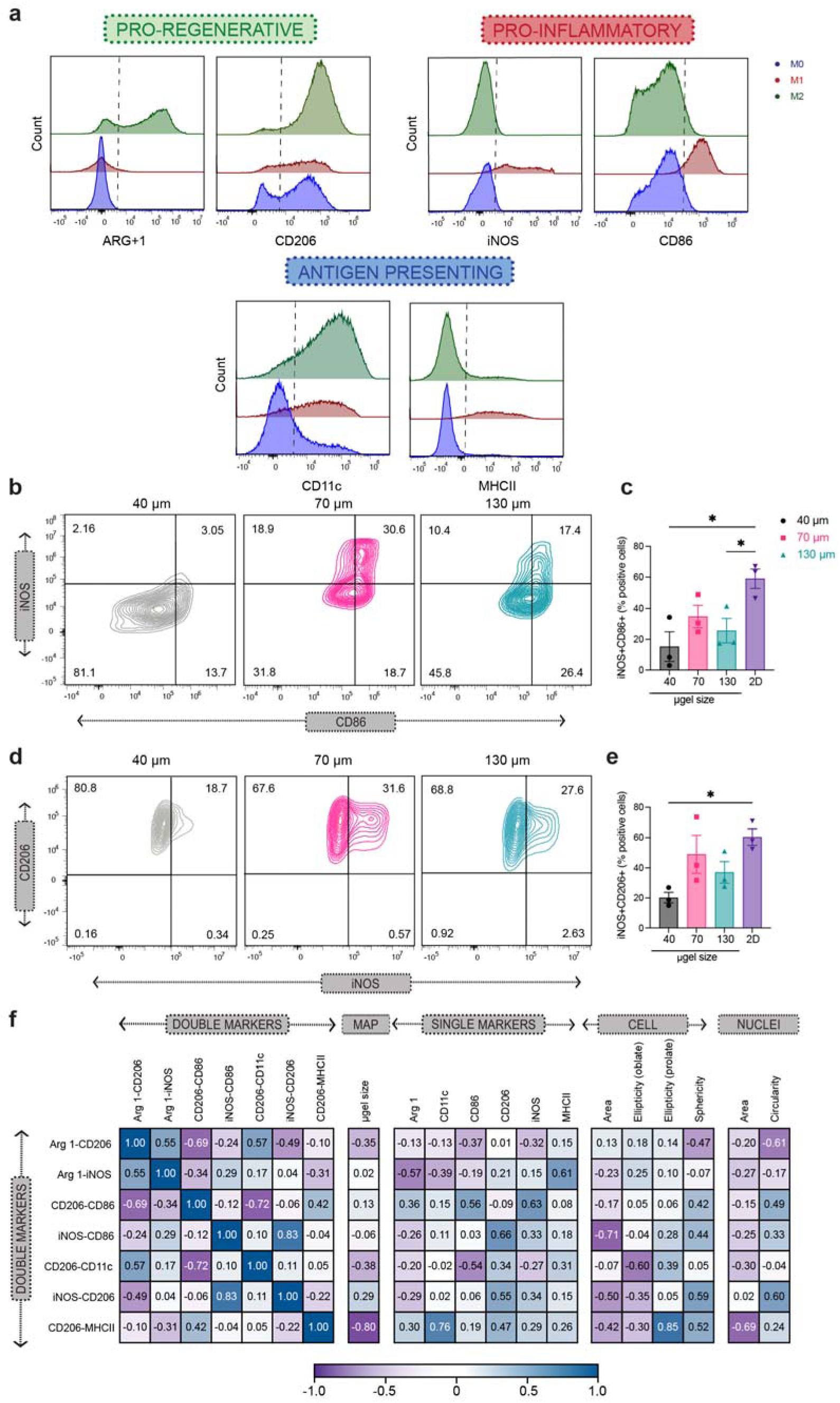
Balance of M1/M2 phenotype was observed upon M1 activation in MAP scaffolds. a, Representative histogram of phenotypical marker expression levels from the flow cytometry analysis of 2D culture BMDM upon M1(LPS/IFNγ) activation, showing pro-inflammatory markers iNOS and CD86, pro-regenerative markers Arg1 and CD206, and antigen-presenting markers CD11c upon M1(LPS/IFNγ) activation. Positive expression threshold was determined as above the MFI of unstimulated BMDM. **b and d,** Representative contour plots of BMDM upon M1(LPS/IFNγ) activation in 40 µm, 70 µm and 130 µm MAP scaffolds, showing iNOS and CD86 co-expression levels as well as iNOS and CD206 co-expression levels. **c and e,** Percentages of double-positive BMDM subpopulation upon M1(LPS/IFNγ) activation for iNOS/CD86 co-expression and iNOS/CD206 co-expression. **f,** Correlation matrix between double-positive BMDM subpopulation upon M1(LPS/IFNγ) activation, selected phenotypical markers, and selected morphological characteristics of BMDM in MAP scaffolds. Statistical analysis: one-way ANOVA with Tukey’s multiple comparisons test made between 40 µm, 70 µm and 130 µm MAP scaffolds groups only when there was significant difference among means. * p<0.05, ** p<0.01, *** p<0.001, **** p<0.0001. Error bars, mean ± s.e.m., n = 3 from three independent experiments per scaffold type.

### Spatial confinement influenced macrophage motility by changing cell trajectory and velocity

Macrophages serve as sentinel cells *in vivo* and are very motile, constantly surveying the our body^3^. We next wanted to visualize macrophages moving in our MAP scaffolds and determine if 3D-pore size affected their movement. When macrophages were encapsulated in a 3D scaffold like MAP scaffolds, they actively explored the void space of MAP scaffolds by moving inside and across 3D-pores and sensing the surface of the microgels (Figure 6a, Supplementary Video 1-3). In 40 µm MAP scaffolds, macrophages wrapped themselves around the small microgels, moving in a circle (Figure 5a, Supplementary Video 1). In 70 µm MAP scaffolds, most macrophages settled into the pocket of 3D-pores and moved from side to side on the surface of microgels (Figure 6a, Supplementary Video 2). Some macrophages in 70 µm MAP scaffolds traveled from one 3D-pore to the adjacent one through the internal doors (Supplementary Video 2). 130 µm MAP scaffolds had the most erratic cell movement. macrophages in 130 µm MAP scaffolds frequently traversed along the surface of the microgels and went through the internal doors into neighboring 3D-pores (Figure 6a, Supplementary Video 3). By tracing cell displacement, we discovered that 130 µm MAP scaffolds posed the least confinement over cell motility whereas macrophages movement in 40 µm and 70 µm MAP scaffolds was limited to the local 3D-pores (Figure 6e). As was show in previous studies, Both cell displacement and velocity were affected by M1/M2 activation^41^, but M1 cytokines elicited the most distinct changes (Supplementary Video 4-9). 130 µm MAP scaffolds had the least restriction over activation-related increase in cell motility, resulting in the largest maximum travel distance and median velocity (Figure 6c,d). The correlation matrix showed a high association between median velocity and maximum distance traveled by the cells with the microgel size of MAP scaffolds and the shape of the cell/nucleus. The bigger 3D-pore size allowed the cells to move and adapt their morphology to the environment freely. In contrast, ellipticity (prolate) in the cell and sphericity in the cell/nucleus showed a negative interaction with the cell motility parameters, indicating that smaller circular cells tended to move less than spread-out cells (Figure 6b). These results suggested that 3D-pore size, imposed by different MAP scaffolds, influenced macrophage motility by changing cell trajectory and velocity.

**Figure 6.**
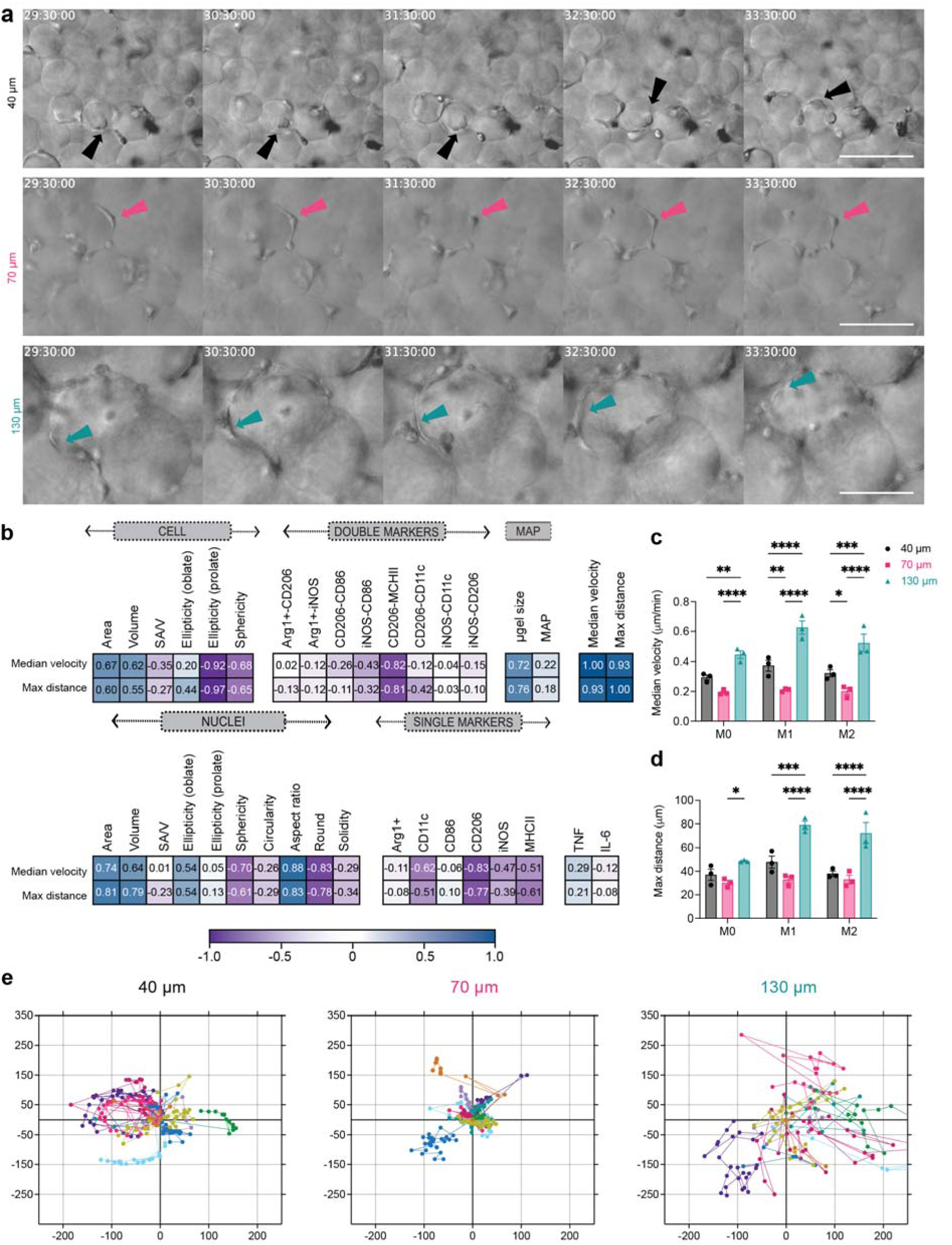
Spatial confinement influenced macrophage motility by changing cell trajectory and velocity. **a,** Representative frames from live-cell imaging video of BMDM upon M1(LPS/IFNγ) activation in 40 µm, 70 µm and 130 µm MAP scaffolds. **b** Correlation matrix between macrophage motility parameters, double-positive BMDM subpopulation upon M1(LPS/IFNγ) activation, phenotypical markers, and morphological characteristics of BMDM in MAP scaffolds. **c,** Median velocity of unstimulated/M0 or M1 or M2 BMDM in MAP scaffolds. **d,** Maximum total travel distance of unstimulated/M0 or M1 or M2 BMDM in MAP scaffolds. **e,** Representative plots of BMDM displacement from origin for 20 hours in 40 µm, 70 µm and 130 µm MAP scaffolds. n = 10 cells in each plot. Statistical analysis: two-way ANOVA with Tukey’s multiple comparisons test made between 40 µm, 70 µm and 130 µm MAP scaffolds groups only when there was a significant difference in scaffold type x treatment. * p<0.05, ** p<0.01, *** p<0.001, **** p<0.0001. Asterisks in black show difference within a treatment and between scaffold type. Error bars, mean ± s.e.m., n = 3 independent experiments per scaffold type.

### Macrophage phenotypes revealed similar patterns between *in vivo* and *in vitro* experiments

Lastly, we characterized the infiltrating macrophages in a subcutaneous implantation model. Each mouse received three injections of 50 µL hydrogel (one of each type 40 µm, 70 µm and 130 µm MAP scaffolds) on the back (Figure 7a). After 7 days post-implantation, the infiltrated cells were extracted, and macrophage phenotypes (CD11b+F4/80+) were analyzed. Four extracellular staining markers, CD86, CD206, MHCII and CD11c, were used to characterize macrophage phenotype (Figure 7b). This time point was chosen to explore the interface between the acute and late inflammatory phase of the foreign body response^37^. The total infiltrated live cell number and CD45+ cell number were similar among all MAP scaffolds groups (Figure 7c and d), which indicated that smaller pore size didn’t pose a significant barrier to cell infiltration. We found that there was a size dependent increase in macrophage percentages, with 130 µm MAP scaffolds having the higher number of macrophages (Figure 7e), possibly due to the larger cell size of macrophages among all immune cells^42^. Similar to our *in vitro* results, 70 µm MAP scaffolds promoted a higher expression of pro-regenerative marker CD206 compared to 40 µm and 130 µm MAP scaffolds (Figure 7f). Pro-inflammatory marker CD86 and antigen-presenting markers MHCII and CD11c had a similar expression among all MAP scaffold groups (Figure 7g-i). Pearson’s correlation matrix comparing our *in vitro* and *in vivo* results showed similar trends for macrophage phenotype, indicating that our *in vitro* results were predictive of the *in vivo* macrophage activation at 7-days (Figure 7l). The general immune response towards all MAP scaffolds was active but constructive, resulting in vascularization in all implants (Figure 7j and m) and an increasing percentage of cellular area mainly in 70 µm MAP scaffold 21-day post-implantation (Figure 7k and m).

**Figure 7.**
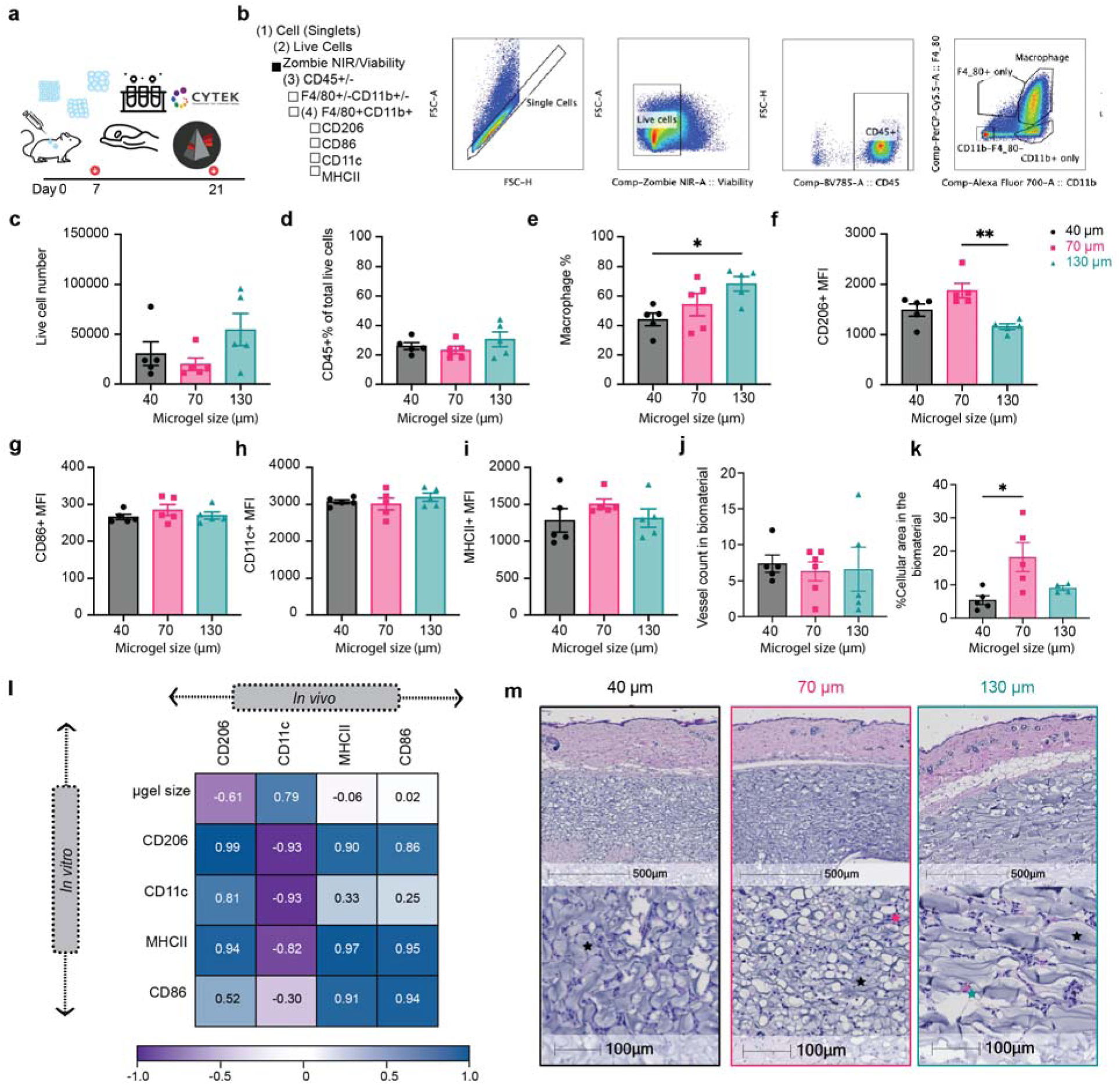
Macrophage phenotypes revealed similar patterns between *in vivo* and *in vitro* experiments. **a**, Scheme illustration of the experiment timeline. After the initial injections, implant extraction and specific assays were performed at day 7 (flow cytometry) and day 21 (H&E)**. b,** Gating strategy for flow cytometry analysis. **c,** The total number of live cells (Zombie NIR-) in different MAP scaffolds groups. **d,** Percentage of CD45+ immune cells (CD45+) in different MAP scaffolds groups. **e,** Macrophages percentage (CD11b+F4/80+) in CD45+ cells in different MAP scaffolds groups. **f-i,** Expression levels of pro-regenerative marker CD206, pro-inflammatory marker CD86 and antigen presenting markers MHCII and CD11c in infiltrating macrophages at day 7. **l,** Correlation plot of the average expression levels of phenotypical markers between *in vivo* and *in vitro* experiments. **j-k,** histologic quantification of vessel count and percentage of granulation tissue in the implant. **m,** Representative pictures of the implants stained with Hematoxylin and eosin (H&E) on day 21. Black stars represent microgel remaining in the implant, pink start represents granulation tissue surrounding the microgels and green-blue star shows vessel formation inside the implant. Statistical analysis: one-way ANOVA with Tukey’s multiple comparisons test made between 40 µm, 70 µm and 130 µm MAP scaffolds groups only when there was significant difference among means. * p<0.05, ** p<0.01, *** p<0.001, **** p<0.0001. Error bars, mean ± s.e.m., n = 5 male mice per scaffold type.

## Conclusions

Macrophages adapt to the features and cues from their surrounding microenvironment. Emerging evidence supports the essential role that physical factors play in modulating macrophage activation. Spatial confinement via pore size and the internal landscape of biomaterials was demonstrated to suppress macrophage LPS-related inflammatory responses in 2D systems^15^. Gaining insights into how spatial confinement regulates macrophage activation in a 3D environment allows us to design more translatable biomaterials for clinical applications.

In this work, we explored the change in macrophage activity when confined within three categories of MAP scaffolds, a granular biomaterial, formed from distinct microgel sizes: 40 µm, 70 µm, and 130 µm diameter. We showed that different degrees of confinement led to different macrophage responses, suggesting that MAP scaffolds are a suitable tool for tuning macrophage behavior. Spatially confining primary mouse macrophages in scaffolds with pore sizes on the scale of cells modulates changes in M1 response, which was associated with a change in cell morphology and motility as well as a reduction of pro-inflammatory markers and cytokines toward a more pro-regenerative antigen-presenting phenotype. Over-confined macrophages in 40 µm MAP scaffolds resulted in smaller cells that stretched to adopt the shape of the smaller 3D-pores. These over-confined cells also showed a higher expression of CD11c, and an increase in the percentage of Arg1+CD206+ macrophage subpopulation, demonstrating the spatially confined macrophages reduced inflammatory response^15^. Confined macrophages in 70 µm MAP scaffolds exhibited more spherical cellular and nuclear shapes. The high sphericity was correlated with higher expression of iNOS, CD206, CD86, and MCHII, representing a macrophage population with a balance of M1/M2 markers. This high expression of CD206 was also observed in a subcutaneous implantation *in vivo* model. In 130 µm MAP scaffolds, which was the least spatially confined scaffold group, cells tended to be larger in both volume and surface area. The higher degree of spatial freedom offered cells more room to stretch and move between 3D-pores, which was seen on live-cell imaging. Similarl to 40 µm MAP scaffolds, we saw a decrease in the expression level of pro-inflammatory markers (iNOS, CD86) in 130 µm MAP scaffolds; however, we also saw lower antigen-presenting marker CD11c in this group. The microgel-size-dependent change in cell morphology correlated with change in nucleus shape, where both 40 µm and 130 µm MAP scaffolds induced a more elongated and less circular nuclei shape. These findings suggested that spatial confinement, imposed by the local size of void space within porous scaffolds, plays a key role in regulating macrophage response in 3D culture and in a subcutaneous implantation model. Future studies that dive deeper into the mechanotransduction pathways governing these relationships in 3D culture will further empower biomaterial design to modulate macrophage response.

## Author contributions

Both YL and ASA made substantial contributions to the design of the work, acquisition, analysis, and interpretation of data. LR provided conceptualization, data and analysis that contributed substantially to figure 1. JX performed part of the experiment and analyzed data that contributed substantially to figures 1.TM performed part of the experiment and analyzed data that contributed substantially to figures 2 and 6. YL, ASA, and TS drafted the manuscript, and all the authors discussed the results and contributed to writing portions of the manuscript and editing the manuscript. TS provided guidance and discussion throughout the project, and made substantial contributions to experimental design, data analysis, and manuscript editing.

## Acknowledgements

We would like to thank the National Institutes of Health (R01AI152568) and all the members of Segura and Collier lab at Duke University for the support. Thank you to our lab alumni Jun Chen, Ph.D., Ethan Ho and our lab member Eleanor Caston for technical assistance in the early stages of this work. Thank you to Duke University Light Microscopy Core Facility (LMCF) and their staff during live imaging acquisition. Thank you to Minerva Matos-Garner, MA from Duke Engineering Graduate Communications and Intercultural Programs for her constructive suggestions on the manuscript and her warm support during the writing process.

## Data availability

The data that support the findings of this study are available from the corresponding authors upon reasonable request. Source data are provided within this paper.

## Methods

### Microgel generation and purification

Microfluidic devices and microgels were produced as previously described. Briefly, a precursor solution with 8-arm PEG Vinylsulfone, K-peptide (Ac-FKGGERCG-NH2, GenScript), Q-peptide (Ac-NQEQVSPLGGERCG-NH2, GenScript) and RGD (Ac-RGDSPGERCG-NH2, GenScript) in 0.3 M triethylamine (Sigma) buffer. The cross-linker solution was prepared by dissolving the di-thiol matrix metalloproteinase-sensitive peptide (Ac-GCRDGPQGIWGQDRCG-NH2, GenScript) GenScript) in distilled water at 12 mM and 10 μM Alexa-Fluor 647-maleimide. These solutions were filtered through a 0.22 um sterile filter before loading them into 1 ml syringes. The final microgels were made of 5% (w/v) 8-arm PEG Vinylsulfone with 500 uM K-peptide, Q-peptide, and RGD, respectively. For small and medium microgel generation, we used a four-inlet device with two inlets for the aqueous solutions (precursor solution and crosslinker solution) and the oil phases (heavy mineral oil with 1% v/v Span-80 as the pinching oil phase and heavy mineral oil with 5% v/v Span-80 and 3% v/v Triethylamine as the gelation oil phase). The microgel solution was collected and allowed to gel overnight at room temperature. For large microgel generation, we used a two-inlet device with one for the aqueous solution (pre-mixed the precursor solution with an equal volume of the crosslinker solution) and the oil phase (heavy mineral oil with 5% v/v Span-80). The microgels were collected in a Span-80 (5% v/v) and Triethylamine (3% v/v) oil bath and allowed to gel overnight at room temperature. These microgels were then purified by repeated washes with a HEPES buffer (0.3 M, pH 8.3 containing 1% Antibiotic-Antimycotic and 2% Pluronic) and centrifugation. The purified microgels were stored in a HEPES buffer (pH 8.3 containing 1% Antibiotic-Antimycotic and 10 mM CaCl_2_) at 4°C. The size of the microgels was measured from microscopic pictures (Nikon C2 confocal microscope, Nikon Instruments Inc.) of three separate batches of microgels using a custom MATLAB code. For cell study, the microgels were swollen in the culture medium for 20 minutes before the study.

### Generation of Scaffold from microgels and Mechanical Testing

Excess buffer from the fully swollen and equilibrated microgels were removed by centrifuging at 22 000 G for 5-20 minutes and discarding the supernatant. 1 μl thrombin (200 U/mL in 200 mM Tris-HCl, 150 mM NaCl, 20 mM CaCl_2_) and 2 μl Factor XIII (250 U/mL) were combined with 50 μl microgels and mixed via thorough pipetting and allowed to incubate at 37°C for 30 minutes to form a solid hydrogel. Storage moduli of the hydrogels were measured by a frequency sweep on rotational rheometry (Anton-Parr, MCR301) at a shear frequency range from 10^-1^ rad/s to 10^2^ rad/s with a strain amplitude of 1%.

### Primary Murine Macrophages culture

Bone marrow-derived macrophages (BMDM) were isolated from 8-12 weeks old C57BL/6 (mix gender) in accordance with institutional and state guidelines and approved by the Duke University’s Division of Laboratory Animal Resources (DLAR) under protocol A019-21-01. Animals were anesthetized, and the tibias and femurs were collected. The bone marrow was flushed out from the bones and broken into cell suspension by repeated pipetting. The cells were differentiated in a culture medium (10% heat-inactivated fetal bovine serum in Iscove’s Modified Dulbecco’s Medium, Gibco) with 15 ng/ml macrophage colony-stimulating factor (Peprotech). The medium was changed on day 1, day 4, and day 8. After full maturation, the cultured cells were tested for murine macrophage pan marker CD11b and F4/80 by flow cytometry to confirm the macrophage percentage (CD11b+F4/80+)^43^.

### Macrophage in vitro encapsulation in MAP Scaffolds

BMDM were detached from the culture flask with TrypLE express enzyme solution (ThermoFisher). The cell pellet was prepared at a final concentration of 10,000 cells per µl of MAP scaffolds. Subsequently, 50 µl of microparticle mixture containing Factor X/Thrombin was added to the cell pellet and thoroughly mixed before pipetting onto a sigmacote-treated glass slide. The slide was then placed into a petri dish and incubated at 37 °C for 45 minutes before transferring to a 6-well plate with the cell culture medium. For imaging purposes, the mixture was injected into the center of a cell culture device made in-house with a coverslip bottom. BMDM were cultured overnight in MAP scaffolds before the 24-hour cytokine activation. M1 activation was with 20 ng/ml LPS (ThermoFisher) and IFN-γ (Peprotech) and M2 activation was with 20 ng/ml IL-4 (Peprotech)^44^.

### Live cell imaging and analysis

10 µl MAP scaffolds with encapsulated BMDM (10,000 cells/µl) were casted in 35 mm dish with 20 mm glass in the bottom (MatTek) and immediately imaged on an Olympus VivaView FL Incubator microscope (20x air objective). After 45 minutes, 3 mL of cell culture medium was added to the dish. The samples were activated with M1/M2 cytokines after 17 hours and imaged for another 27 hours. Images were taken at an interval of 30 minutes. For analysis, at the 24-hour mark, 10 cells were chosen at random. These 10 cells were tracked using the manual tracking feature in Fiji software (ImageJ), which allows the user to mark the location of each cell in every frame. If a cell disappeared at any point, tracking was stopped to ensure all data points could be attributed to only one cell. The data collected for three videos per scaffold were used to create the plots of cell displacement and determined the median velocity and max distance traveled by the cells in each scaffold.

### IHC staining

At the designated time points, the samples were fixed with 4% PFA overnight at 4°C or 15 minutes at room temperature and then permeabilized with 0.3% Triton X-100 in 1xPBS for 10 minutes at room temperature. This was followed by staining for F-actin via Alexa Fluor 488/rhodamine phalloidin (Life Technologies) overnight at 4 °C. The scaffolds were then washed with 1X PBS, followed by counterstaining with a DAPI solution in 1X PBS for 15 minutes at room temperature. The samples were imaged with a Nikon C2 confocal microscope (Nikon Instruments Inc.), using a 40x magnification water immersion lens. The height of image stacks was set at 130 µm and the total number of slices was 260.

### Cell morphology analysis

To analyze cell morphologies within the scaffolds, cell renderings were created of the 3D confocal images (40x objective) using the IMARIS x64 software (Oxford Instruments). To complete these renderings, both “spots” and “surfaces” functions were used. First, a computer-generated plot was created in the DAPI channel to visualize the cell nuclei and 9 μm was chosen as an estimation of the nuclei size. By overlaying the original image on top of the generated spots, background signals and nuclei without any actin filament around them were removed from the plot. A similar process was used with the “surfaces” function to view the morphology of the cells. The renderings were completed with the surface detail set at 1 μm. The absolute intensity threshold was adjusted manually to ensure that the surface matched the actin filaments in the actual images. A baseline of absolute intensity threshold was chosen for each scaffold type and minor adjustments were made to avoid over- or under-rendering. Cell clusters were defined as any surface containing two or more nuclei and were therefore analyzed as one unit. Cell surfaces without a nucleus in close proximity were removed manually. For all the surfaces created, a list of parameters was generated for further analysis: sphericity, ellipticity, volume and area of a single cell and a cell cluster, total cell count, cluster number, and single cell number.

### Nuclei morphology analysis

To analyze nuclear shape within the scaffolds, mean Z-stack projection images were generated using Fiji software (ImageJ). After splitting and thresholding in the DAPI channel with “Intermodes” method, the binary image was processed with “fill holes” and “watershed” functions. Four parameters were calculated: nuclei area, circularity (defined as “4*pi(area/perimeter^2^)”), roundness (defined as “4*area/pi*sqr(major axis)”), and aspect ratio (defined as “major axis/minor axis”).

### Flow cytometry study

At the designated time points, the cells were extracted from MAP scaffolds gels by enzymatic digestion of the gel with digestion solution (200 U/ml Collagenase IV and 125U/ml DNase I) in RPMI media for 10 minutes at 37 °C. Similarly, for in vivo studies the implants were extracted and diced finely prior to enzymatic digestion with the digestion solution (200 U/ml Collagenase IV and 125U/ml DNase I) in RPMI media for 15 minutes at 37°C. The resulting material was filtered through a 70 μm cell strainer and washed once with 1xPBS to get a single-cell suspension. These cells were then stained with a Zombie NIR (BioLegend) solution for 15 minutes at room temperature and blocked with Fcr Blocking Reagent (Miltenyi Biotec) for 10 minutes on ice, followed by surface marker staining for 30 minutes on ice. For intracellular marker staining, an intracellular fixation & permeabilization buffer set (Thermo Fisher) was used to prepare the samples (Supplementary Table 1). After staining, samples were washed and resuspended in 150 ul flow buffer (1x PBS, 1 mM EDTA, 0.2% BSA, 0.025% proclin) and analyzed on the Cytek NL-3000 Flow Cytometer. Data was acquired using SpectroFlo software and analyzed using FlowJo 8 (TreeStar) flow cytometry data analysis software. The relative abundance of the macrophage population was gated following the gating strategy. Relative abundance of macrophage subpopulations was determined as a fraction of CD11b+F4/80+ live cells (gated first on scatter FSCxSCC, then doublet discrimination via FSC-AxFSC-H, prior to Viability dye).

### ELISA

Cell culture medium was collected from 2D and MAP scaffolds confined BMDMs after 24 h of cytokine stimulation. To remove cell debris, the cell culture medium was centrifuged, and the supernatant was collected and stored at −80 °C until further processing. All ELISA kits were purchased from ThermoFisher Scientific, and the tests were performed according to the manufacturer’s protocol.

### Subcutaneous implantation

7-12-week-old male C57BL/6 mice (Jackson Laboratory) were anesthetized with 3.0% isoflurane and maintained at 1.5-2.0% isoflurane. The microgels and crosslinker solution were thoroughly mixed and loaded in a 1 cc syringe with a 29-gauge needle. Each mouse received three injections of 50 µL hydrogel (one of each 40 µm, 70 µm and 130 µm MAP scaffolds) on the back. After injection, mice were monitored until full recovery from the anesthesia. All procedures were approved by the Duke University Institutional Animal Care and Use Committee and followed the NIH Guide for the Care and Use of Laboratory Animals.

### Histology staining

At day 21, the implants were extracted for histology examination. For paraffin embedding, samples were fixed with 4% paraformaldehyde overnight at 4 °C and further processing. Paraffin blocks were sectioned into 5 μm thickness with at least 3 serial-sections per slide for hematoxylin and eosin (H&E) staining. The sections were de-waxed and hydrated using xylene then decreasing ethanol concentrations. They were stained in Mayer Hematoxylin Solution (EMS) for 15 minutes before being rinsed in warm running tap water for 15 minutes. They were placed in DI water for 30 seconds, 95% ethanol for 30 seconds, and then into Alcoholic Eosin Y Counterstain (EMS) for 30 seconds. They were then dehydrated and cleared before being mounted in DPX (EMS).

### Computational experiments

Granular scaffolds comprising monodisperse spheres were simulated with SideFX Houdini software using their rigid-body physics solver. Spherical particles were dropped through a funnel into a 600 x 600 x 600 µm3 container to achieve random packing. Particle diameter was adjusted according to experimental groups, while all other parameters were held constant. Simulated scaffold data of particle centers and radii were then inputted into in-house software, termed LOVAMAP, for void space analysis. LOVAMAP uses particle configuration and Euclidean distance transforms of the void space to extract subtypes of the medial axis that are used to segment the space into 3D-pores. We then compute and report the volume of 3D-pores, and we focus our computational analysis on interior pores that do not extend to the outside of the scaffold in order to avoid edge effects. For lab-derived MAP scaffolds, we first convert confocal z-stack images of each scaffold into a data format that lists the 3D-voxels associated with each unique particle, then run the data through LOVAMAP. We report the volume of all 3D-pores for our lab-derived MAP scaffolds since the limitations of our microscope’s depth of field results in few interior 3D-pores relative to simulated data.

### Statistical analysis

All experiments were performed in three biological replicates with three separate primary cell extractions from mouse bone marrow, and three technical replicates per biological replicate. Two-way ANOVA was used to establish the significant of microgel size and activation type. In Figure 2 b-h we combined unstimulated, M1 and M2 groups (3 points per biological replica) in each MAP GELS condition because the two-way ANOVA test indicated that the activation condition was not a significant factor. For *in vivo* experiments 5 mice per time point were used as biological replicas. One-way ANOVA coupled with post hoc Tukey test was performed to compare all variables against microgel size, considering significant p-values of less than 0.05. To analyze the interaction between multiples variables, Pearson correlation plots were used where values higher than ±0.5 meant moderate to high correlation^45^ between three independent biological replicas. Analysis and plots were made using GraphPad Prism software 9 (GraphPad Software).

